# Branched chain amino acids modulate the proteomic profile of Pro-induced differentiation for the infective stages of *Trypanosoma cruzi*

**DOI:** 10.1101/2024.04.17.589966

**Authors:** Janaina de Freitas Nascimento, Flávia Silva Damasceno, Sabrina Marsicobettre, Francisca Natália de Luna Vitorino, Renan Weege Achijan, Julia Pinheiro Chagas da Cunha, Ariel Mariano Silber

## Abstract

*Trypanosoma cruzi*, the causative agent of Chagas disease, undergoes a complex life cycle involving triatomine insects as vectors and mammals. The differentiation of epimastigote forms into metacyclic trypomastigotes within the insect vector is crucial for the parasite’s life cycle progression. Factors influencing this process, including temperature, pH, and nutritional stress, along with specific metabolite availability, play a pivotal role. Amino acids like Pro, His, and Gln support cell differentiation, while branched-chain amino acids (BCAAs) inhibit it. Interestingly, combining the pro-metacyclogenic amino acid Pro with one of the anti-metacyclogenic BCAAs results in viable metacyclics with significantly reduced infectivity. To explore the characteristics of metacyclic parasites differentiated in the presence of BCAAs, proteomics analyses were conducted. Metacyclics obtained in triatomine artificial urine (TAU) supplemented with Pro alone and in combination with Leu, Ile, or Val were compared. The analyses revealed differential regulation of 40 proteins in TAU-Pro-Leu, 131 in TAU-Pro-Ile, and 179 in TAU-Pro-Val, as compared to metacyclics from TAU-Pro. Among these, 22%, 11%, and 13% of the proteins were associated with metabolic processes, respectively. Notably, enzymes related to glycolysis and the tricarboxylic acid (TCA) cycle were reduced in metacyclics with Pro-BCAAs, while enzymes involved in amino acid and purine metabolic pathways were increased. Furthermore, metacyclics with Pro-Ile and Pro-Val exhibited elevated enzymes linked to lipid and redox metabolism. These findings suggest that the presence of BCAAs can reshape the metabolism of metacyclics, contributing to the observed reduction in infectivity in these parasites.

## Introduction

*Trypanosoma cruzi* is the causative agent of Chagas disease, a neglected disease endemic of the Americas and affecting worldwide 6-7 million people [1]. *T. cruzi* has a complex life cycle occurring among triatomine insect vectors and mammals. Inside the triatomines, the parasites proliferate as non-infective epimastigotes, which colonize the digestive tube. In the terminal portion of the digestive tube, epimastigotes differentiate into the infective, non-proliferating forms metacyclic trypomastigotes, which are responsible for initiating the infection in mammals, including humans. The differentiation process from epimastigotes to metacyclic trypomastigotes is known as metacyclogenesis. This process occurs naturally at the rectal ampoule of the triatomine [reviewed by [2]], and can be reproduced *in vitro* by using the chemically defined medium denominated triatomine artificial urine (TAU), which mimics the physicochemical characteristics of the insect urine [3]. TAU medium must be supplemented with nutrients that are able to support metacyclogenesis after a starvation period [3, 4]. The most commonly used formulations are the ones supplemented with a combination of glucose (Glc) and the amino acids proline (Pro), glutamate (Glu) and aspartate (Asp) (TAU 3AAG) [3]. However, it has been shown that some amino acids can support metacyclogenesis *in vitro* by themselves [4, 5]. It is tempting to attribute this metacyclogenic activities of metabolites such as glucose, Pro, Asp, Glu and glutamine (Gln) to their demonstrated role as carbon and energy sources. However, not every amino acid fueling ATP production is able to support metacyclogenesis: epimastigotes do not differentiate in the presence of the branched-chain amino acids (BCAAs) leucine (Leu), isoleucine (Ile) and valine (Val) as the only carbon source in TAU [3, 4]. Even more, when TAU-Pro, is supplemented with any of the BCAAs metacyclogenesis is significantly diminished [4], and very interestingly, the diminished population of metacyclics that can be obtained in the presence of BCAAs are less infective [6]. Thus, regardless of their role in ATP production, some amino acids can support metacyclogenesis, acting as ‘pro-metacyclogenic’ and others somehow inhibit this process, acting as ‘anti-metacyclogenic’.

Pro can induce metacyclogenesis in different *T. cruzi* strains such as Dm 28, Dm 30, CL, Perú isolate, CL Brener and CL14 [3, 4, 6], and the participation in this process of each enzyme involved in Pro oxidation has been analyzed [7]. In addition to this, Pro also plays an important role in many other biological processes in *T. cruzi*, such as energy metabolism and serving as a carbon source [8–10], intracellular differentiation [11], host-cell invasion [12], resistance to thermal and nutritional stress [13] oxidative imbalance [9, 13, 14] and osmotic control [15]. In order to be available for the many functions it is involved in, Pro is obtained by *T. cruzi* from the extracellular environment [16] and is able to biosynthesize it from Glu [17, 18]. The oxidation pathway of Pro to Glu occurs in two enzymatic and one non-enzymatic steps: first proline dehydrogenase (ProDH) converts Pro into Δ^1^-pyrroline-5-carboxylate (P5C) [9]. P5C undergoes a non-enzymatical chemical reaction by hydrolyzing the pyrrolic ring, rendering glutamate-γ-semialdehyde (GSA), which is the substrate of the P5C dehydrogenase (P5CDH), which finally oxidizes it into Glu [8]. In yeast, P5CDH can be inhibited by Leu and Ile, [19] and in *T. cruzi* Leu at high concentrations also inhibits P5CDH [6].

Given that: (i) Pro is a ‘pro-metacyclogenic’ amino acid; (ii) BCAAs do not support the differentiation from epimastigote to trypomastigote metacyclic; (iii) BCAAs impair metacyclogenesis supported by Pro and (iv) metacyclics obtained in the presence of BCAAs are less infective, we analyzed the global proteomic profiles of the metacyclic trypomastigotes obtained from TAU medium supplemented with different metabolic substrates: 3AAG, Pro, Pro-Leu, Pro-Ile and Pro-Val. Globally, we found that parasites differentiated in TAU-3AAG and TAU-Pro medium are very similar, whereas the comparison between metacyclics from TAU-Pro versus TAU-Pro-BCAAs showed changes in the proteomic profile, mostly in proteins involved in metabolic processes, translation, proteolysis and regulation of gene expression.

## Experimental Procedures

### *Trypanosoma* strains and culture

Epimastigotes from CL strain clone 14 were maintained in the exponential growth phase by subculturing every 48 h in LIT medium [20] supplemented with 10% FCS at 28 °C.

### Metacyclogenesis and metacyclic trypomastigotes purification

Stationary phase epimastigotes were obtained from a starting exponential cell culture at 5 × 10^6^ parasites/mL, kept for 4 days in LIT at 28°C. Parasites were washed with PBS (137 mM NaCl; 2.6 mM KCl; 8 mM Na_2_HPO_4_; 1.4 mM KH_2_PO_2_, pH 7.4) at 1,600 x *g* for 5 min, transferred to TAU (190 mM NaCl; 17 mM KCl; 2 mM MgCl_2_; 2 mM CaCl_2_; 8 mM potassium phosphate buffer, pH 6.0) at a final concentration of 5 x 10^7^ parasites/mL and incubated for two hours at 28 °C. After, parasites were harvested and transferred to one of the TAU media of interest: TAU 3AAG (TAU + 10 mM Glc; 10 mM Asp; 10 mM Glu; 10 mM Pro; pH 6.0), TAU Pro (TAU + 10 mM Pro), TAU Pro-Leu (TAU Pro + 10 mM Leu), TAU Pro-Ile (TAU Pro + 10 mM Ile), TAU Pro-Val (TAU Pro + 10 mM Val) at a final concentration of 5 x 10^7^ parasites/mL. After 6 days of incubation, the differentiation rate was estimated by counting the parasites in the Neubauer chamber. Then, parasites were washed twice with PBS, pH 8.0 and the metacyclic trypomastigotes forms were isolated by ion exchange chromatography on DEAE-cellulose (Sigma) as previously described [21].

### Protein Extraction and Digestion

Approximately 1 x 10^8^ purified metacyclic trypomastigotes obtained in each differentiation condition were used to prepare protein extracts. Cells were resuspended in 250 µL lysis buffer (8 M urea, 75 mM NaCl, 50 mM Tris, pH 8.2) supplemented with protease / phosphatase / deacetylase inhibitors: 2 mg/L leupeptin, 1 μM pepstatin and 10 mM pyrophosphate, 0.1 mM sodium orthovanadate, 0.1 mM PMSF, 1 mM glycerophosphate, 5 mM sodium butyrate, 1 mM NaF. After protein quantification, 60 μg of proteins were added to 100 mM Tris-HCl pH 8.5. Next, trichloroacetic acid (TCA) was added to a final concentration of 20%. Samples were incubated for 16 h at 4°C and centrifuged at 14,000 x *g* for 30 min at 4°C. Finally, pellets were washed twice with 500 μL of cold acetone (14,000 x *g* for 10 min) and kept at room temperature to dry. The pellets from precipitation were resuspended in 30 μL of lysis buffer (8 M urea, 75 mM NaCl, 50 mM Tris pH 8.2) and reduced by adding 5 mM Dithiothreitol (DTT) for 25 min at 56°C. Alkylation was performed by adding 14 mM iodoacetamide for 30 min at room temperature in the dark. Next, 5 mM of DTT was added, and samples were incubated for another 15 min at room temperature in the dark. The urea concentration was reduced with 25 mM Tris pH 8.2 and 100 mM CaCl_2_. Finally, trypsin (1 μg/μL; Sigma-Aldrich) was added in the proportion of 1:200 (enzyme:substrate) for 16 h at 37°C under agitation. After incubation, the reaction was stopped with 1 μL of 100 % trifluoroacetic acid (TFA) and dried in speed-vac. Digested peptides were desalinized using the Sep-Pak® Light tC18 Cartridges (Waters, Milford, MS, USA) columns. First, the columns were conditioned with 100% methanol, followed by a solution of 0.1% TFA and 50% acetonitrile (ACN) and a solution of 0.1% TFA. Next, samples were resuspended with 0.1% TFA and loaded to the column. After washing the columns with 0.1% TFA, samples were eluted with 0.1% TFA/50% ACN solution, dried in the speed-vac, and analyzed by mass spectrometry.

### Mass Spectrometry-Based Analysis of *T. cruzi* Tryptic Peptides

Digested peptides from three biological replicates were resuspended in 0.1% formic acid and fractionated on an in-house reverse-phase capillary column (10 cm × 75 µm, filled with 5 µm particle diameter C18 Aqua resins-Phenomenex) coupled to nano-HPLC (NanoLC; ThermoFisher Scientific). The fractionation was carried out through at a 60 min gradient of 5-35% gradient ACN in 0.1% formic acid followed for 30 min gradient of 35-95% at a flow rate of 200 nl/min. The eluted peptides were analyzed online on the LTQ-Orbitrap Velos High-Resolution Mass Spectrometer (ThermoFisher Scientific) (source voltage of 1.9 kV, capillary temperature of 200°C). The mass spectrometer was operated in DDA mode with dynamic exclusion enabled (exclusion duration of 45 s), MS1 resolution of 30,000, and MS2 normalized collision energy of 35. For each cycle, one full MS1 scan range of 200–2,000 m/z was followed by ten MS2 scans (for the most intense ions) using an isolation window size of 2.0 m/z and collision-induced dissociation (CID).

### Analysis and integration of proteomic data

Data from LTQ Velos-Orbitrap were analyzed using the MaxQuant (v. 1.6.1) software using the TriTrypDB database (taxonomy DB32 T. *cruzi* CL Brener Esmeraldo like and Non-Like haplotype). Parameters were set with tolerance in the MS of 4.5 ppm and MS/MS of 0.5 Da, carbamidomethylation of cysteine as a fixed modification: oxidation of methionine and acetylation of protein N-termini as variable modifications; and a 1% False Discovery Rate (FDR). The match between runs option was used to increase the number of trusted IDs. Normalized LFQ values of the proteingroups.txt output were used to obtain the quantitative values. Next, Perseus (v. 1.5.6.0 and 1.6.2.1) program (from the MaxQuant package) identified differentially expressed proteins and integrated the data with available public information. Those proteins identified as potential false positives (contaminants, reversals, and identified by only one site) were removed. Finally, statistical tests (T-test with FDR permutation) were used to identify proteins differentially expressed between conditions, accepting only reproducible results (p <0.05; s0: 0.1). The mass spectrometry proteomics data have been deposited to the ProteomeXchange Consortium via the PRIDE partner repository (PXD049190).

### Bioinformatic Analysis

For cluster analysis by the Fuzzy c-Means (FCM) algorithm, average LFQ values for each protein of each analyzed condition were used to generate clusters using http://computproteomics.bmb.sdu.dk/Apps/FuzzyClust/ [22]. Gene Ontology Biological Process and Cell Component analysis were performed using the tools available on TriTrypDB (https://tritrypdb.org/tritrypdb/) and data were manually curated. Venn diagrams were generated using InteractiVenn (https:www.interactivenn.net/) [23]. The PCA decomposition was performed using MATLAB’s “pca” function with default parameters (that is, performing data centering, attributing the same weight to all data points and using singular value decomposition as the underlying algorithm).

### Western blotting

Western blotting was used to determine the expression levels of tyrosine aminotransferase (TAT) in total cell lysates of metacyclic trypomastigotes. Briefly, parasites were harvested as described above and resuspended in lysis buffer (20 mM Tris-HCl pH 7.9; 0.25 M sucrose; 1 mM EDTA pH 8.0; 0.1% (v/v) Triton X-100; 1 mM PMSF; 2 μg/mL aprotinin; 0.1 mg/mL TLCK and 10 μM E-64). Samples were sonicated on ice and clarified by centrifugation (10,000 x *g* for 30 min at 4 °C). Protein concentration was determined by the Bradford method using bovine serum albumin (BSA) as a standard [24]. Equal amounts of protein samples (30 μg) were loaded per lane and submitted to electrophoresis (SDS-PAGE) in gels containing trichloroethanol (Sigma-Aldrich). Proteins were transferred into 0.2 μm nitrocellulose membranes (BioRad), blocked in PBS-0.1% Tween-20 (PBST) supplemented with 5% (w/v) skimmed milk powder and probed (1 h at RT) against specific sera. The enzyme TAT was probed with a polyclonal rabbit antiserum (1:500), gently gifted by Prof. Cristina Nowicki, University of Buenos Aires [25]. The fluorescence signal of the total protein for each extract contained in the membranes after UV light exposure was used as the loading control. Membranes were washed three times and incubated with anti-mouse IgG or anti-rabbit IgG HRP-linked antibodies (Cell Signaling Technology) both diluted in PBST (1:2,000). The chemiluminescence reaction was performed using SuperSignal West Pico Chemiluminescent ECL substrate (Thermo Scientific) following the manufacturer’s instructions.

### Pyruvate kinase activity assay

The activity of pyruvate kinase (PK) was determined in total cell lysates of metacyclic trypomastigotes obtained as previously mentioned. Equal amounts of protein extracts of metacyclic trypomastigotes obtained in TAU Pro or TAU Pro-BCAAs (50 μg) were used for each reaction. The reaction mixture contained: 78 mM potassium phosphate buffer pH 7.6; 1.16 mM phospho(enol)pyruvic acid (PEP) (Sigma-Aldrich); 0.22 mM NADH disodium salt trihydrate (Amresco); 13.6 mM MgSO_4_; 3 mM ADP sodium salt (Sigma-Aldrich); 10 U L-lactic dehydrogenase (Sigma-Aldrich). The reactions were monitored by spectrophotometry using a Spectramax i3 plate reader (Molecular Devices), at 28 °C, in 96 wells plates following the decrease in absorbance at 340 nm due to NADH oxidation in the coupled lactate dehydrogenase reaction. For each reading, the same reaction mixture without PEP was used as control/blank. The media and standard deviation of two biological replicates were represented in a graph using GraphPad Prism 10 software.

## Results

### Metacyclogenesis in the presence of different amino acids induces distinct protein expression profiles

To investigate the protein expression profiles of the metacyclic trypomastigotes differentiated in the presence of different metabolites, we induced metacyclogenesis in TAU supplemented with either 3AAG, Pro, Pro-Leu, Pro-Ile, and Pro-Val (Figure S1). Metacyclic trypomastigotes were purified and their proteome profile was evaluated by LC-MS/MS. After initial analysis, we identified 1,457 proteins as present in all samples (Table S1). Principal component analysis showed that the proteomic profile of the metacyclics obtained in TAU-3AAG and TAU-Pro cluster together, whilst the metacyclics differentiated in the presence of Pro-BCAAs produce distinct proteomic profiles (Figure 1). Fuzzy c-means clustering [22] of LFQ values grouped approximately 28% of the identified proteins in 7 distinct clusters, evidencing different trends of protein expression in each particular differentiation condition (Table S2). To look in more detail at the proteins present in each cluster, we analysed the Gene Ontology annotation of biological process and cell compartment for each protein (Figure 2). The expression of proteins in Clusters 1 and 4 changed in response to the presence of either Pro-Ile or Pro-Val during cell differentiation. Approximately 18% of the proteins more expressed in these conditions (Cluster 1) participate in metabolic processes, mostly in amino acids metabolism, including tyrosine aminotransferase (TAT) which is responsible for the transamination step of the BCAAs degradation pathway [26]. Accordingly, proteins in Cluster 1 appear to be, aside from cytoplasmic, localised in mitochondria, glycosomes and the contractile vacuole. In contrast, less expressed proteins in metacyclics differentiated in TAU Pro-Ile and TAU Pro-Val (Cluster 4) are involved in oxidation-reduction, other metabolic processes, such as carbohydrates and purine metabolic processes, and post-transcriptional regulation of gene expression (Figure 2A, Table S3 and S6). Proteins involved in oxidation-reduction processes also appear less expressed in metacyclics differentiated in the presence of Pro-Leu (Cluster 2). Surprisingly, there is a high incidence of retrotransposon hot spot proteins (RHSPs) and proteins localised as integral components of membranes in Cluster 2 when compared to other clusters. On the other hand, proteins involved in post-transcriptional regulation of gene expression as well as in translation and RNA processing are up-regulated in metacyclics differentiated in TAU Pro-Leu (Cluster 3) (Figure 2B, Tables S4 and S5). Cluster 5 group proteins which expression reduces in cells differentiated in TAU Pro-Val. These are mostly involved in translation, protein folding and post-transcriptional regulation of gene expression, and include ribosomal proteins, translation initiation factors (eIFs) and RNA-binding proteins (RBPs). Correspondingly, these proteins mostly localise to the cytoplasm, nucleus, axoneme and ribosomes. In turn, proteins involved in proteolysis and other metabolic processes such as carbohydrates and fatty acids metabolic processes are up-regulated in response to the presence of Pro-Val during metacyclogenesis and localise to the mitochondria, glycosomes and contractile vacuoles, apart from the cytoplasm (Figure 2C, Tables S7 and S9). Interestingly, cluster 6 shows proteins with increased expression in TAU Pro-Ile, which mostly take part in translation, oxidation-reduction and amino acid metabolic processes, but there is no correspondent mirroring cluster (Figure 2D, Table S8). Together, the data suggest that, although the BCAAs similarly inhibit metacyclogenesis supported by Pro, they induce different proteomic profiles on the resulting trypomastigote metacyclics. Noteworthy, all clusters also point to a very similar trend of protein expression when we compare trypanosomes differentiated in TAU 3AAG and TAU Pro.

**Figure 1:**
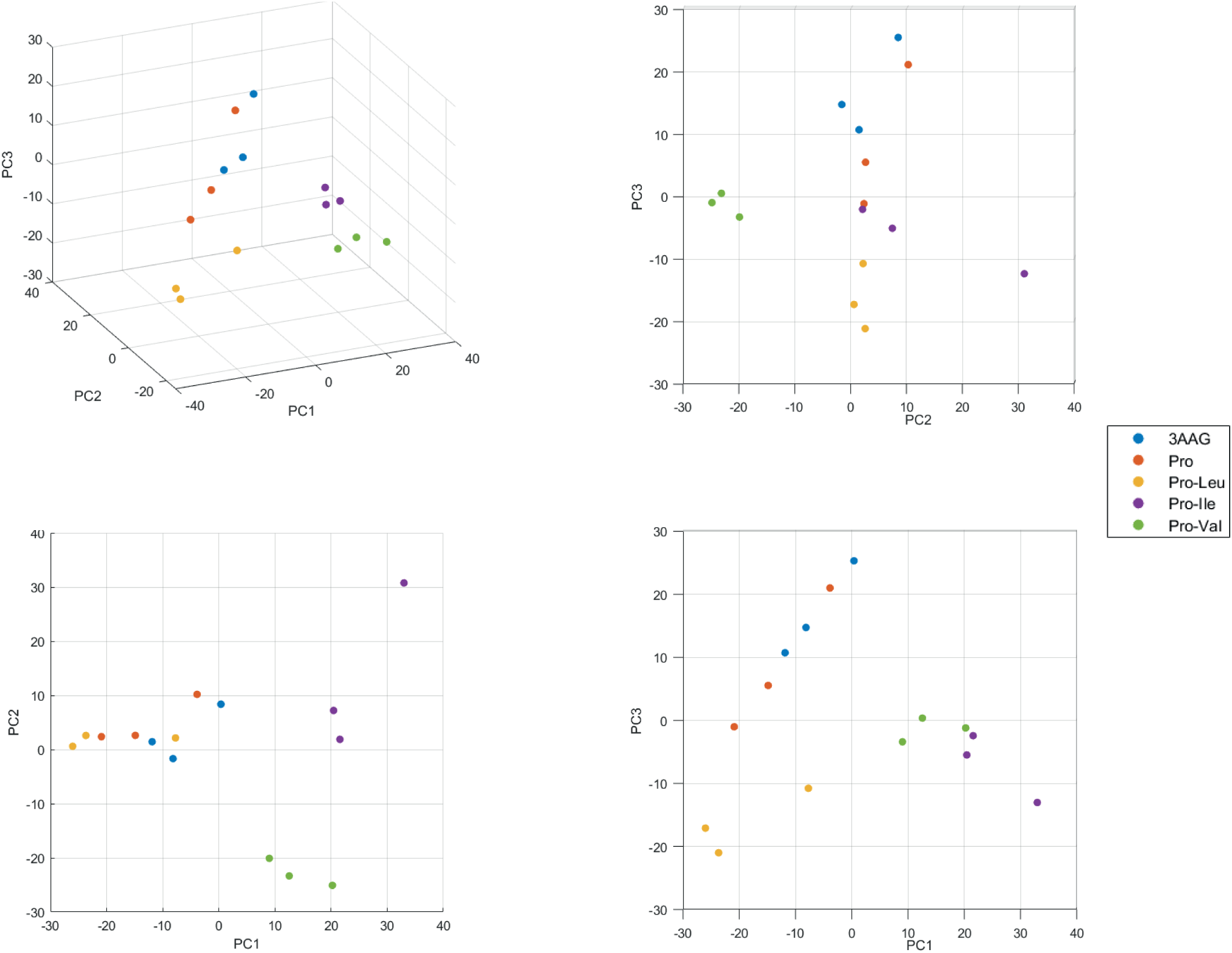
Distribution of the experimental data in principal component space, seen as a 3D view of the first three principal components, and projections on the planes defined by those principal components taken in pairs. The variability explained by PC 1, 2 and 3 are, respectively, 27%, 16% and 14%. The other PCs explain 8% or less of the variability each. The points corresponding to 3AAG (blue) and Pro (purple) cluster together, with Pro-Leu (yellow) in its immediate vicinity, while Pro-Ile (orange) and Pro-Val (green) are further apart, both from the (3AAG, Pro, Pro-Leu) cluster and from each other. Each circle corresponds to an independent biological replicate for each differentiation condition.

**Figure 2:**
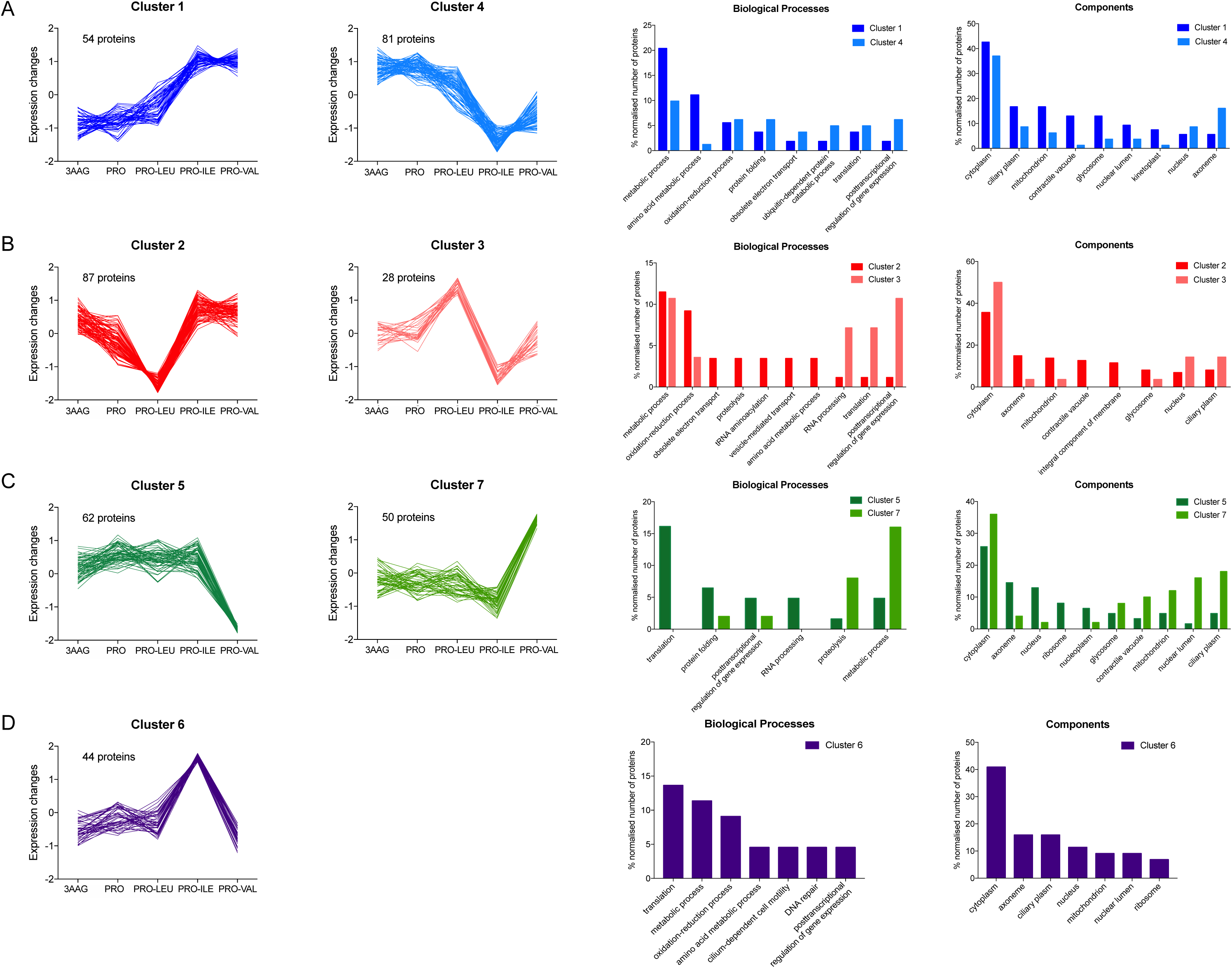
Fuzzy c-means clustering analysis shows distinct protein expression profiles in metacyclics differentiated in TAU 3AAG, Pro, Pro-Leu, Pro-Ile and Pro-Val. Clusters evidence variations in the proteomic profiles of metacyclics differentiated in **A.** TAU Pro-Ile and Pro-Val; **B.** TAU Pro-Leu; **C.** TAU Pro-Val and **D.** TAU Pro-Ile. Grouped bar graph shows the distribution of the proteins of each cluster according to biological process and cell components. Average LFQ values were used for clustering analysis. Computed and Curated GO Biological Processes available in TriTrypsDB were used to generate bar graphs.

### Metacyclogenesis in TAU 3AAG and TAU Pro induce similar proteomic profiles

*In vitro* metacyclogenesis performed in TAU 3AAG and TAU Pro produce similar differentiation rates [27]. In order to evaluate whether metacyclic trypomastigotes differentiated in the presence of 3 amino acids plus glucose (TAU 3AAG) or just Pro (TAU Pro) have also similar protein expression profiles, we performed differential expression analysis. Metacyclic trypomastigotes obtained in TAU Pro presented a very similar proteomic profile when compared with metacyclics differentiated in TAU 3AAG. A few proteins are differentially expressed between these two differentiation conditions: 39 proteins in total, from which 17 are up-regulated and 22 are down-regulated in parasites differentiated in TAU Pro (Figure S2, Tables S10 and S11). Around 35% of the up-regulated proteins are annotated as hypothetical. Gene Ontology analysis evidence that most of these proteins are localized to cytoplasm, glycosome, nucleus, ribosome and axoneme. Up-regulated proteins are predominantly involved in energy production, nucleotide production, DNA organization and ribosome composition. On the other hand, of the 22 down-regulated proteins in TAU Pro parasites, only 18% are identified as hypothetical. Down-regulated proteins are predominantly related to the cytoplasm or nucleus and are mostly involved in energy metabolism and regulation of gene expression, respectively. Interestingly, proteins involved in metabolism such as proline racemase [28] and malate dehydrogenase [29] are downregulated in these parasites (Tables S12 and S13).

### The relationship between BCAAs and Pro

Considering the importance of Pro in metacyclogenesis and the negative interference of BCAAs, we compared the proteomic profile of parasites differentiated in TAU Pro medium with parasites differentiated in TAU Pro supplemented with each one of the BCCAs. We then excluded the proteins present in the differential expression analysis between metacyclic trypomastigotes from TAU 3AAG and TAU Pro because they are likely to be a result of the presence of other amino acids and glucose in the TAU 3AAG. Since different subsets of proteins are differentially expressed in response to the presence of each BCAA during metacyclogenesis, we describe the results of each comparison separately below.

### Comparison of proteomic patterns of metacyclic trypomastigotes from TAU Pro with TAU Pro-BCAAs

Leu causes the strongest reduction in the Pro-induced metacyclogenesis (see [4, 6] and Figure S1). However, differential expression analysis comparing the proteomic profiles of metacyclic trypomastigotes obtained in TAU Pro and TAU Pro-Leu shows only 47 differentially regulated proteins, from which 11 proteins were excluded for being present in the comparison between metacyclics differentiated in TAU 3AAG and TAU Pro. From the remaining proteins, 20 are down-regulated and 16 are up-regulated in parasites differentiated in TAU Pro-Leu (Figure 3A, Tables S14 and S15). Approximately 35% and 25% of the proteins are annotated as hypothetical among the down and up-regulated datasets, respectively. In order to identify the biological processes in which the differentially expressed proteins are possibly involved, Gene Ontology analysis was performed (Figure 3B-C, Tables S16 and S17). The results evidenced that down-regulated proteins are involved in biological processes such as the generation of energy, posttranscriptional regulation of gene expression and oxidation-reduction processes; on the other hand, up-regulated proteins participate in respiration and translation processes. In terms of cell components, down-regulated proteins are enriched in the cytoplasm and mitochondria when compared to the up-regulated proteins that are mostly present in axoneme, contractile vacuole and ribosomes.

**Figure 3:**
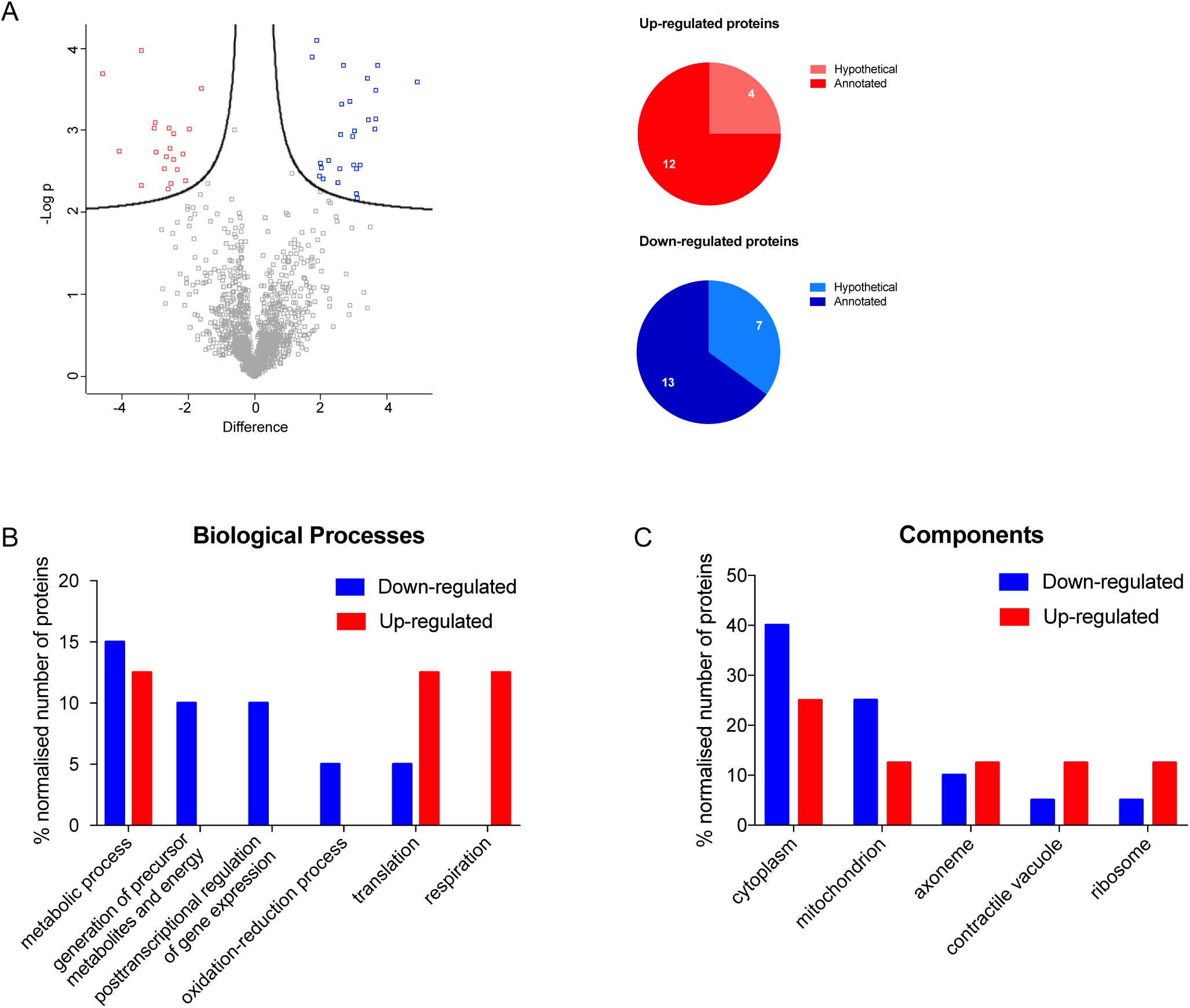
Parasites differentiated in TAU Pro-Leu have a similar proteomic profile to those differentiated in TAU Pro. **A.** Volcano plot of LFQ values of proteins from metacyclic trypomastigotes differentiated in TAU Pro and TAU Pro-Leu. Differentially expressed proteins are shown in blue (down-regulated in TAU Pro-Leu) and red (up-regulated in TAU Pro-Leu). Pie charts show the number of hypothetical proteins in the sets of up and down-regulated proteins. **B** and **C.** Distribution of up and down-regulated proteins according to biological process and cell component, respectively.

The next step was to analyze the metabolic changes induced in metacyclic trypomastigotes by the addition of Ile to the TAU Pro-defined medium. Differential protein expression analysis comparing the proteomic profile of metacyclics obtained in TAU Pro and TAU Pro-Ile showed 154 differentially regulated proteins, from which 23 proteins were excluded for being present in the comparison between metacyclics differentiated in TAU 3AAG and TAU Pro. The remaining 131 proteins are divided into 68 down-regulated and 63 up-regulated proteins. Among the differentially expressed proteins, approximately 35% of the down-regulated proteins are annotated as hypothetical, whilst only 20% of the up-regulated proteins have the same annotation (Figure 4A, Tables S18 and S19). Gene Ontology analysis showed that proteins putatively involved in translation, post-transcriptional regulation of gene expression and protein folding are down-regulated in response to the presence of Pro-Ile during cell differentiation, whereas proteins involved in ubiquitin-dependent protein catabolic process and amino acid metabolic processes are up-regulated. However, there are no major differences in the distribution of up and down-regulated proteins in the cell compartments (Figure 4B and 4C, Tables S20 and S21). Noteworthy, there are possible changes in the BCAAs degradation pathway in parasites differentiated in TAU Pro-Ile, with up-regulation of tyrosine aminotransferase (TAT) [26] and down-regulation of the beta subunit of BCKDH complex, which performs the second step of the pathway [30].

**Figure 4:**
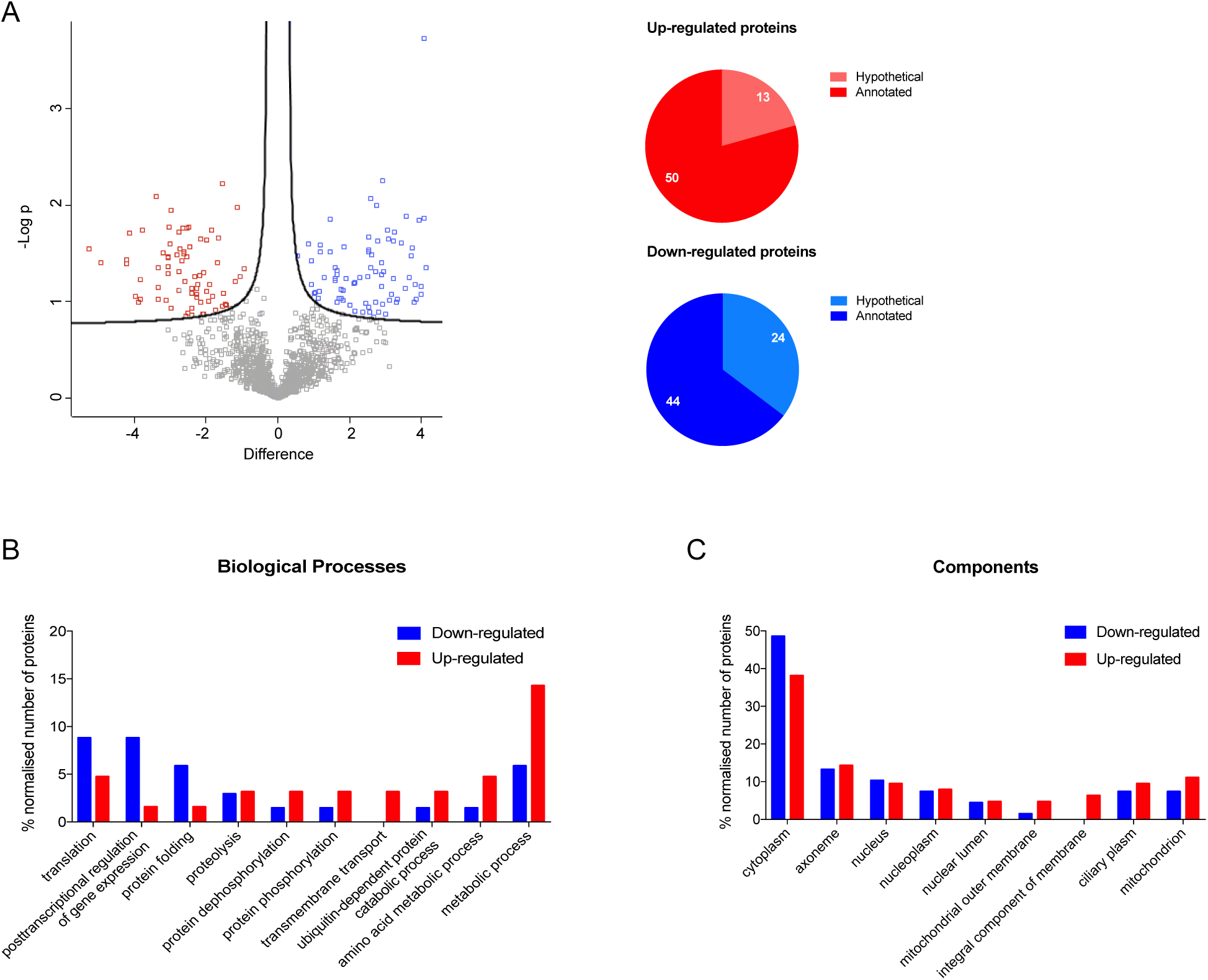
Addition of Ile to TAU Pro induces proteomic changes in the resulting metacyclics. **A.** Volcano plot of LFQ values of proteins from metacyclic trypomastigotes differentiated in TAU Pro and TAU Pro-Leu. Differentially expressed proteins are shown in blue (down-regulated in TAU Pro-Ile) and red (up-regulated in TAU Pro-Ile). Pie charts show the number of hypothetical proteins in the sets of up and down-regulated proteins**. B** and **C.** Distribution of up and down-regulated proteins according to biological process and cell component, respectively.

In contrast with the comparisons previously described, differential expression analysis between metacyclic trypomastigotes obtained in TAU Pro and TAU Pro-Val showed the highest number of differentially regulated proteins, 197 in total, from which 18 proteins were excluded for being present in the comparison between metacyclics differentiated in TAU 3AAG and TAU Pro. Among the remaining proteins, 78 are down-regulated and 101 are up-regulated (Figure 5A, Tables S22 and S23). Approximately 30% of the down-regulated proteins are annotated as hypothetical whereas only 20% of the up-regulated proteins have the same annotation. Gene Ontology analysis showed down-regulation of proteins involved in biosynthetic processes (such as translation and protein folding) and up-regulation of proteins involved in metabolic processes (such as oxidation-reduction processes and proteolysis) (Figure 5B, Tables S24 and S25).

**Figure 5:**
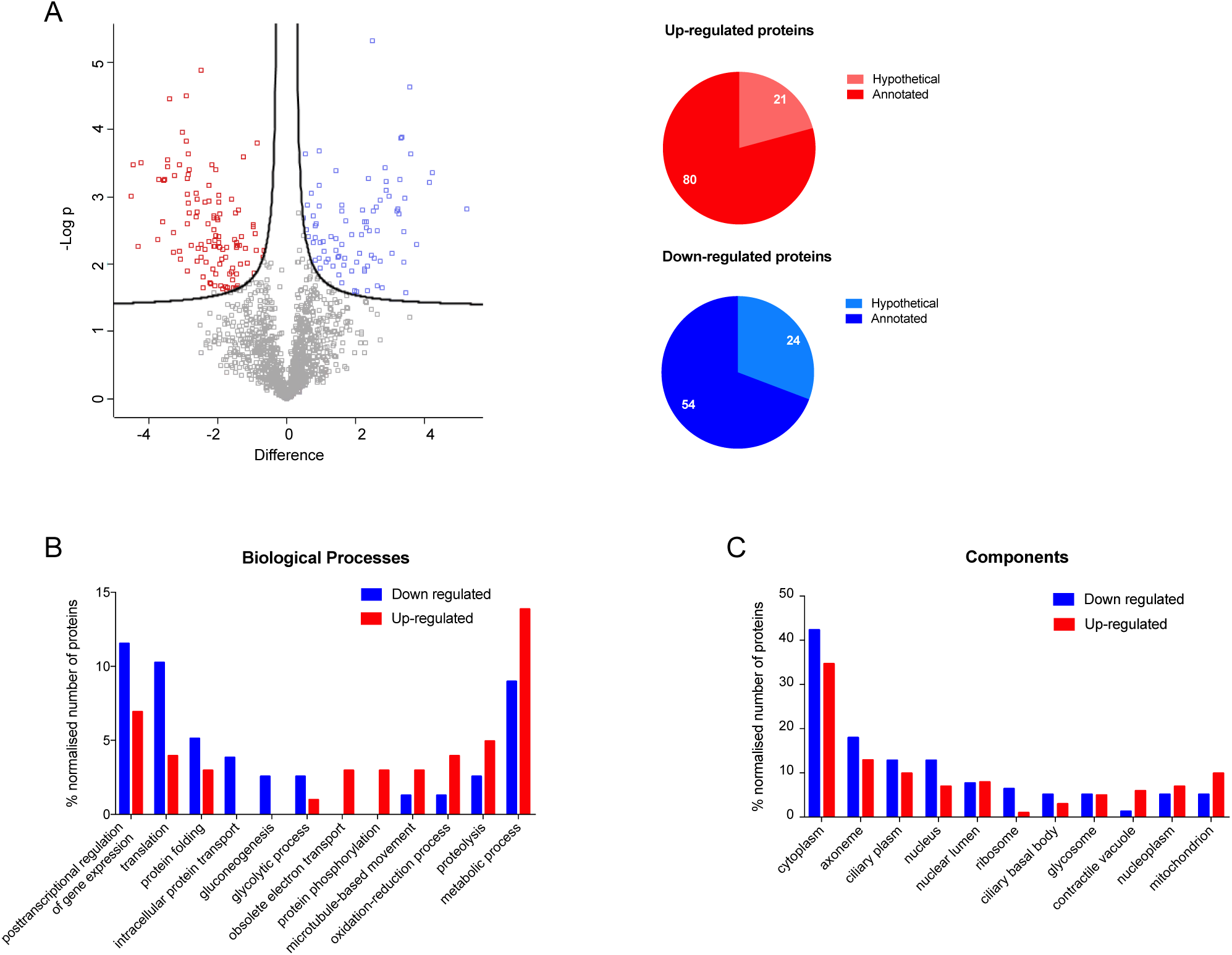
Metacyclogenesis in TAU Pro-Val results in changes in proteins related to metabolic processes. **A.** Volcano plot of LFQ values of proteins from metacyclic trypomastigotes differentiated in TAU Pro and TAU Pro-Leu. Differentially expressed proteins are shown in blue (down-regulated in TAU Pro-Val) and red (up-regulated in TAU Pro-Val). Pie charts show the number of hypothetical proteins in the sets of up and down-regulated proteins. **B** and **C.** Distribution of up and down-regulated proteins according to to biological process and cell component, respectively.

Related to metabolic processes, results suggest that there are changes in Glu, Pro, Ser, Thr and Gln metabolism. Unlike the previous conditions, there are changes in glucose metabolism. The enzymes phosphoglycerate kinase (PGK) [31], 2,3-bisphosphoglycerate-independent phosphoglycerate mutase (iPGAM), glycerol kinase (GK) and fructose-1,6-bisphosphatase (FBPase) that are involved in glycolysis and/or gluconeogenesis are differentially regulated. There is also differential regulation of two putative subunits of the respiratory chain – down-regulation of ubiquinol-cytochrome c reductase (UCYT CR) and up-regulation of cytochrome c oxidase VIII (COXVIII), also the putative ATP synthase is down-regulated, which might lead to alterations in energy production in these parasites. In the BCAAs degradation pathway, there is up-regulation of TAT, responsible for the first step of the pathway. Interestingly, proteins related to fatty acid metabolism are up-regulated, such as fatty acyl-CoA synthetase 2 and carnitine O-palmitoyltransferase II, both putative. Moreover, proteins associated with the structure and motility of the cell are differentially expressed in metacyclics differentiated in TAU Pro-Val, such as the putative microtubule-associated protein Gb4 and flagellum attachment zone 4 which are down-regulated and the putative cytoskeleton-associated protein, flagellar member 1 and a trypanosoma basal body component are up-regulated.

### Final comparison TAU Pro x TAU Pro-BCAAs

Protein expression levels of over 300 proteins varied in response to the presence of Pro combined with the different BCAAs during metacyclogenesis. However, only a few proteins are similarly differentially regulated when we compare the datasets of up and down-regulated proteins of metacyclic trypomastigotes differentiated in TAU Pro-Leu, Pro-Ile and Pro-Val (Figure 6). Interestingly, the autophagy-related protein 24 is up-regulated in metacyclics differentiated in the presence of TAU Pro supplemented with any of the BCAAs, whereas the flagellum attachment zone protein 4 (FAZ4) is down-regulated in those parasites (Tables S26 and S27). Moreover, proline dehydrogenase, glutathione-S-transferase and fatty acyl-CoA synthetase are up-regulated in TAU Pro-Ile and TAU Pro-Val metacyclics. Different trans-sialidases are down-regulated in TAU Pro-Leu and TAU Pro-Val when compared with TAU Pro. Furthermore, the enzyme tyrosine aminotransferase (TAT) is up-regulated in parasites differentiated in TAU Pro-Ile and TAU Pro-Val, whereas pyruvate kinase (PK) is down-regulated in parasites differentiated in TAU Pro-Ile when compared with TAU Pro. PK activity and TAT expression were analysed in the crude extract of metacyclics differentiated in TAU Pro and TAU Pro+BCAAs (Figure S3). The results confirm the differential expression of PK and TAT identified in the proteomic analysis. Taken together, the data indicate that metacyclogenesis in the presence of each BCAA induces differences in the proteomic profiles of the metacyclic trypomastigotes.

**Figure 6:**
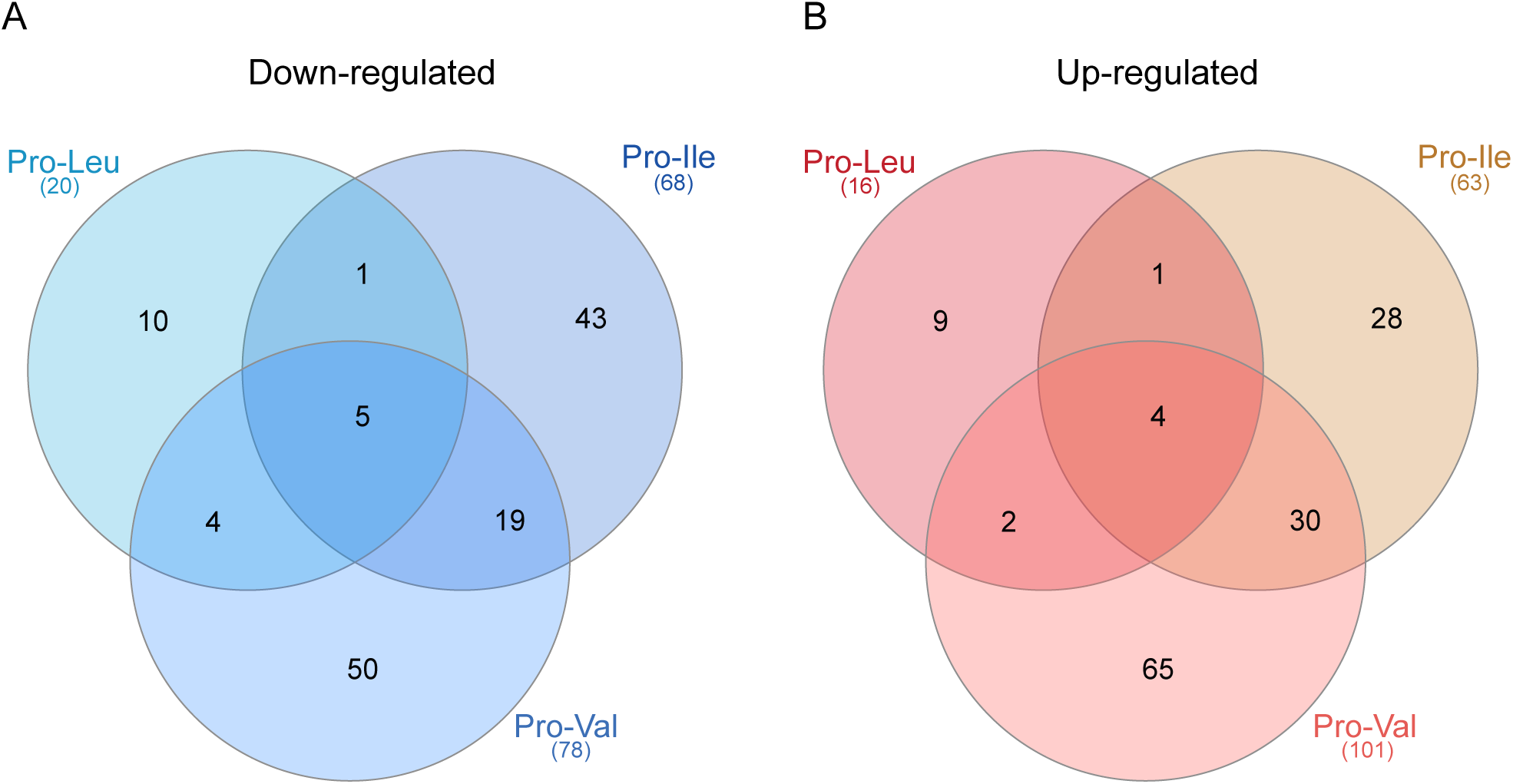
Each of the BCAAs induces particular proteomic changes in the metacyclic trypomastigotes. Venn diagrams shows the number of proteins shared between the datasets of down (A) and up-regulated (B) proteins of metacyclics differentiated in TAU Pro-Leu, Pro-Ile and Pro-Val when compared to TAU Pro.

## Discussion

Metacyclogenesis occurs at the final portion of the triatomine’s gut. In this adverse environment, the parasite faces intense nutritional stress, which has been shown to trigger cell adhesion and metacyclogenesis [3, 32]. Early studies have shown the presence of several amino acids in the excreta of the insect vector *Rhodnius prolixus*, such as histidine, glycine, phenylalanine, Leu and Val [33]. More recently, metabolomic analysis of faeces of different triatomine species showed that phenol lipids and amino acids are the most abundant components of the insect’s excreta [34].

Other factors have been shown to impact the metacyclogenesis rate either *in vivo* and/or *in vitro,* such as the trypanosome redox state [35], the temperature in the insect vector [36] and the regulation of autophagy [37, 38]. One other important factor which determines the success of cell differentiation is the exposure to different substrates [3, 5, 39, 40]. Some substrates such as the amino acids Pro, Asn and Gln are ‘pro-metacyclogenic’, supporting cell differentiation. Other substrates such as the branched-chain amino acids (BCAAs) Leu, Ile and Val are ‘anti-metacyclogenic’ not only because they do not support metacyclogenesis but also inhibit the action of the metabolites that do so. As it has been previously shown, the presence of Pro combined with any of the BCAAs during *in vitro* metacyclogenesis reduces cell differentiation rates [4, 6]. In this work, we investigated if the metacyclic trypomastigotes originated in these conditions present important differences in their protein expression profiles.

Proteomic analysis showed that the proteomic profile between parasites differentiated in TAU 3AAG and TAU Pro medium is very similar, which could be expected since Pro is one of the components of TAU 3AAG [27]. However, the comparison between the proteomic profiles of the metacyclics differentiated in TAU Pro versus TAU Pro supplemented with each BCAA presented more than three hundred proteins differentially expressed, suggesting that BCAAs might not only interfere directly in the Pro metabolism during metacyclogenesis but also in other biological processes.

Metacyclogenesis induced in TAU Pro supplemented with Leu, Ile or Val resulted in lower differentiation rates when compared to metacyclogenesis induced in TAU Pro. Interestingly, parasites differentiated in TAU Pro and TAU Pro-Leu showed the most similar proteomic profile whilst parasites obtained in the presence of Pro-Val showed the most significantly different proteomic profile. Maybe more strikingly is the fact the different proteins involved in a variety of cell processes are differentially regulated in each specific condition, evidencing that despite the similar phenotype in the reduction of Pro-induced metacyclogenesis, each BCAA distinctly affects metacyclogenesis. Some of the biological processes that might be affected will be discussed in more detail because their relevance in the biology of *T. cruzi* has already been demonstrated.

*T. cruzi* uses BCAAs as a carbon and energy source and their catabolism produces intermediates for the tricarboxylic acid cycle [40, 41]. The canonic enzyme BCAA transaminase is absent in this parasite; the first step of oxidation of BCAAs is catalyzed by TAT and aspartate aminotransferase (ASAT) [26, 42]. These non-canonical aminotransferases differ in their catalytic competencies towards BCAAs; TAT seems to more readily utilize Leu, while ASAT appears to more actively transaminate Ile [26]. Considering that TAT is almost exclusively expressed in epimastigotes and that ASAT is expressed in negligible levels in this stage but that its abundance notably increases in the mammalian stages of this protozoan, the difference in substrate preference might reflect specific nutritional needs throughout the life stages of this pathogen [26]. Once the BCAAs are transaminated, their carbon skeletons undergo different metabolic fates. Theoretically, Leu degradation leads to the formation of the ketogenic metabolites acetoacetate and acetyl-CoA, whilst Ile degradation generates acetyl-CoA and propionyl-CoA which are ketogenic and glucogenic compounds. Finally, Val degradation forms succinyl-CoA which is glucogenic [43]. In trypanosomatids, Leu can be used as a direct precursor and carbon source for sterol and isoprenoid biosynthesis and its chain carbons are also utilized for fatty acid production, whereas Ile is poorly used for sterol synthesis and, when used in an anabolic pathway, is more efficiently channelled to fatty acid biosynthesis [44, 45]. In addition, it was reported that Glc causes an increment in the metabolism of Leu to CO_2_ production, revealing a connection between Leu metabolism and glycolysis [46]. Leu metabolism diverges with respect to Ile and Val by the availability of a specialized acyl-CoA dehydrogenase: isovaleryl-CoA dehydrogenase (IVDH); actually, this fact has been studied in our lab. The differences in the metabolic fates of the BCAAs might explain the phenotypic particularities observed in the metacyclic trypomastigotes differentiated in the presence of these amino acids.

Transamination of BCAAs has been shown to play a key role in α-ketoglutarate, Glu and NH_3_ homeostasis in different types of cells [47–49]. Interestingly TAT was found to be up-regulated in metacyclic trypomastigotes differentiated in TAU Pro-Ile and TAU Pro-Val, but not in TAU Pro-Leu. Additionally, other enzymes putatively involved in pyruvate and Glu metabolism are differentially expressed in metacyclics obtained in the presence of the BCAAs, such as the down-regulation of glutamate dehydrogenase [50] in TAU Pro-Leu and up-regulation in TAU Pro-Ile, down-regulation of pyruvate kinase 2 [51] in TAU Pro-Ile and up-regulation of glutamine synthetase [52] in TAU Pro-Val. Glutamine synthetase (GS) has been described as an ammonium detoxifier in *T. cruzi* amastigotes [52] so it is reasonable to speculate that GS up-regulation occurs in response to ammonium concentration resulting from the increased amino acid catabolism in parasites differentiated in the presence of Pro+BCAAs.

Atwood et al., identified proteins from different stages of *T. cruzi* by LC-MS/MS and concluded that the differentiation into metacyclic trypomastigotes resulted in an increased expression of proteins involved in antioxidant defenses [53]. Our results indicate the regulation of proteins involved in oxidation-reduction in response to the presence of BCAAs during the differentiation process, confirming the importance of this process for this form of the parasite. Polyamines and thiol-containing molecules have also been shown to be involved in the maintenance of the redox balance of *T. cruzi* [54], which in turn plays a major role in the success of the mammalian host infection [55]. More recently, the metabolism of polyamines has also been implicated as a positive regulator of metacyclogenesis in *T. cruzi* [38]. Our results indicate a differential expression of proteins involved in glutathione metabolism, such as glutathione synthetase [56], which is down-regulated in parasites differentiated in the presence of Pro-Ile, whereas a putative glutathione S-transferase is up-regulated in metacyclics differentiated in TAU Pro-Ile and TAU Pro-Val. The data point out that the presence of BCAAs in the medium led to possible differential regulation of polyamine biosynthesis in metacyclic trypomastigotes.

During nutritional stress and metacyclogenesis, translation is attenuated [57], *T. cruzi* uses proteins previously accumulated in reservosomes as amino acid sources [58] and, accordingly, inhibition of proteasome activity prevents metacyclogenesis [59]. Interestingly, proteins putatively involved in proteolysis such as proteasome activator protein pa26 [60] and the putative proteasome regulatory ATPase subunit 1 are up-regulated in metacyclics differentiated in TAU Pro-Ile and TAU Pro-Val whereas several proteins involved in translation, including ribosomal proteins and eukaryotic initiation factors (eIFs) are down-regulated in these parasites when compared to metacyclics differentiated in TAU Pro. However, this relationship seems to be the opposite in parasites differentiated in TAU Pro-Leu when we compare with metacyclics obtained in TAU Pro and TAU 3AAG (Clusters 2 and 3).

The flagellum of invasive trypomastigotes plays an important role in the infection process, driving the trypanosome to initiate and disseminate the infection in the body [61]. Functional analysis of flagellar proteins has now revealed surprising new roles for the flagellum in trypanosomatids, such as cell morphogenesis, cell division, and immune evasion [62]. Our results show that proteins involved in the flagellum structure are differentially regulated in metacyclic trypomastigotes differentiated in TAU Pro supplemented with Ile or Val. Proteins such as inner arm dynein 5-1, dynein heavy chain and paraxonemal rod are up-regulated in metacyclics differentiated in TAU Pro-Val; only dynein light chain was up-regulated in TAU Pro-Ile derivate trypomastigotes. On the other hand, radial spoke protein 3, C-terminal motor kinesin, outer arm dynein and microtubule-associated protein Gb4 are down-regulated in TAU Pro-Val trypomastigotes and only radial spoke 3 is downregulated in TAU Pro-Ile. Dyneins are described as important for the structure and motility of eukaryotic cilia and flagella [63, 64]. The outer and inner arm dyneins bound to the A-tubule of doublet microtubules, walk on the neighbouring B-tubule and provide the driving force for motility [62, 65]. Outer arm dyneins are generally uniform in composition and may contain either two or three distinct heavy chains, depending on the organism [66]. In *T. brucei*, the knockdown of the outer dynein arm by RNAi led to the absence of propulsive waves and loss of forward motility [67]. Some assembled protein modules such as radial spokes, nexin links and central pair projections are important for axonemal function through their roles as cross-linkers and regulators of doublet microtubules sliding and bending. [65]. Radial spokes provide a platform for the assembly of signaling proteins and are part of a mechanochemical signal transduction system that regulates dyneins [62, 68, 69]. Radial spoke 3 has been investigated in *T. brucei* and it is required for parasite motility [67, 70]. These proteins contain predicted regulatory domains, supporting the concept that spokes play important roles in signal transduction [70] and are modifiers of dynein activity and flagellar beating [62]. The flagellar attachment zone (FAZ) is an adhesion region of *T. cruzi* epimastigote form where the flagellum emerges from the flagellar pocket and remains attached to the cell body [71]. Studies have suggested that the FAZ plays a role in cellular organization and cytokinesis; FAZ structures are replicated and associated with the new flagellum [72]. Also, flagellum attachment zone protein 4 (FAZ4) is downregulated in all conditions in which the BCAAs were added, which might affect the capability of cell invasion of the parasite, once that is important to parasite motility [73]. The proteins involved in the structure of the flagellum are important for this parasite stage, however, the fact that these proteins are mostly differentially regulated in cells differentiated in TAU Pro-Ile and mostly in TAU Pro-Val needs to be more investigated.

Trans-sialidases are cell surface proteins responsible for the incorporation of sialic acid from host cells into molecules present in the parasite membrane and play a role in immune evasion and host cell entry mechanisms. Trans-sialidases, TS-like and mucin members are likely to be the most abundant proteins on the surface of *T. cruzi* [74]. This family of proteins display greater variability; proteins of group II that have no trans-sialidase activity are capable of binding to β-galactose, laminin, fibronectin, collagen, cytokeratin and are involved in cell adhesion and invasion [75]. By proteomic analysis, De Godoy et al, report that all of the 18 trans-sialidases identified were up-regulated in metacyclic trypomastigotes when compared with the initial phases of differentiation of *T. cruzi* [76]. Cordero et al reported that in metacyclic forms 39% of the total expressed proteins are surface proteins (members of the trans-sialidase superfamily, surface glycoproteins, mucin and GP63 protease) and there are involved in host-parasite interplay confirmed that this developmental form expresses a large repertoire of surface glycoproteins involved in the host-cell adhesion and invasion [77]. In our case, some specific putative trans-sialidases proteins are downregulated in trypomastigotes obtained in TAU Pro-Leu and TAU Pro-Val when compared to metacyclics obtained in TAU Pro. Together, differential regulation of flagellar proteins and trans-sialidases might affect the infection capability of these parasites, but this needs to be further investigated.

When we analysed the global proteomic profiles of the metacyclics differentiated in the presence of the BCAAs, we found that only a few proteins were similarly differentially expressed between the conditions. One of the proteins that is up-regulated in all three conditions is an autophagy-related protein, ATG24. Autophagy has been reported as one of the main processes that regulate metacyclogenesis in *T. cruzi* [37, 38] and the ortholog of ATG24 in *T. brucei* has been associated with the inhibition of cell differentiation [78], what indicate that differentiation in the presence of BCAAs might be interfering with the regulation of autophagy.

## Concluding remarks

In this work, we analysed the proteomic profile of trypomastigote metacyclics differentiated in the TAU medium supplemented with different substrates. The result demonstrates that the presence of different metabolites during metacyclogenesis can induce changes in the proteomic profiles of the parasites. Parasites that differentiate in the presence of Pro, Glu, Asp and Glc (3AAG medium) present a more similar proteomic profile than the parasites that differentiate in Pro only, while the parasites differentiate in Pro medium supplemented with each BCAA present a different proteomic profile when compared with parasites differentiated in the presence of Pro. The presence of Pro combined with either Leu, Ile or Val in the differentiation medium changes the expression of proteins related to metabolic processes such as amino acid metabolism, as well as oxidation-reduction and translation, mainly. Taken together these data show that the metabolites present in the differentiation medium can modify protein expression profiles which can determine the differentiation rates and might affect the infectivity of the parasites.

## Acknowledgments

This work was supported by: Fundação de Amparo à Pesquisa do Estado de São Paulo (FAPESP) 2021/12938-0 (awarded to AMS), and 2018/15553-9 (awarded to JCPC) Conselho Nacional de Pesquisas Científicas e Tecnológicas (CNPq) grant 307487/2021-0 (awarded to AMS) and Wellcome Trust grant 222986/Z/21/Z (awarded to JFN and AMS).

## Data availability

The mass spectrometry proteomics data have been deposited to the ProteomeXchange Consortium via the PRIDE partner repository (PXD049190).

## Supplemental data

This article contains supplemental data: Supplemental Figures S1-S3 and Supplemental Tables S1-S27.

## Abbreviations

3AAG: Three amino acids and glucose
ACN: Acetonitrile
ASAT: Aspartate aminotransferase
Asp: Aspartate
BCAAs: Branched-chain amino acids
BCKDH: Branched-chain α-ketoacid dehydrogenase complex
BSA: Bovine serum albumin
COXVIII: Cytochrome c oxidase VIII
eIFs: eukaryotic initiation factors
FAZ4: Flagellum attachment zone protein 4
FBPase: Fructose-1,6-bisphosphatase
FCM: Fuzzy c-Means
FCS: Fetal calf serum
FDR.: False Discovery Rate
GK: Glycerol kinase
Glc: Glucose
Gln: Glutamine
Glu: Glutamate
GS: Glutamine synthetase
GSA: glutamate-γ-semialdehyde
Ile: Isoleucine
iPGAM: 2,3-bisphosphoglycerate-independent phosphoglycerate mutase
Leu: Leucine
LIT: Liver infusion tryptose
P5C: Δ1-pyrroline-5-carboxylate
P5CDH: P5C dehydrogenase
PCA: Principal component analysis
PEP: phospho(enol)pyruvic acid
PGK: Phosphoglycerate kinase
PK: Pyruvate kinase
Pro: Proline
ProDH: Proline dehydrogenase
RHSPs: Retrotransposon hot spot proteins
Ser: Serine
TAT: Tyrosine aminotransferase
TAU: Triatomine artificial urine
TCA: Trichloroacetic acid
Thr: Threonine
UCYT: CR Ubiquinol-cytochrome c reductase
Val: Valine

